# Functional trait differences between invasive and non-invasive *Lonicera* (Caprifoliaceae) species

**DOI:** 10.64898/2026.11.13.766506

**Authors:** Isha Saini, Olivia Thorgersen, Michael LaScaleia

## Abstract

Functional traits can influence a plant’s ability to establish and spread in a new environment. Examining traits associated with resource acquisition and defense may help explain why some introduced species become invasive while others do not. This study aimed to determine whether invasive and non-invasive *Lonicera* species differed in growth-related, physical-defense, and chemical-defense traits at the Arnold Arboretum of Harvard University in Boston, Massachusetts. Invasive species were predicted to exhibit traits associated with greater resource acquisition while maintaining distinct, but not necessarily reduced, chemical defenses. To investigate these differences, 12 traits were assessed across six invasive and eight non-invasive species, with the non-invasive group comprising three native and five non-invasive exotic species. Overall, trait composition differed between invasive and non-invasive species. Invasive species exhibited higher specific leaf area and lower leaf thickness, suggesting that invasiveness among the sampled species was associated with differences in leaf resource acquisition and structure, rather than a consistent reduction in chemical defense. Future research should include additional individuals and populations to determine whether these patterns occur more broadly and can help predict plant invasiveness.

## INTRODUCTION

Across the world, invasive exotic plant species are becoming increasingly widespread and harmful to native ecosystems (Pyšek et al. 2020). Consequently, a key area of current ecological research aims to determine what functional traits promote plant invasiveness (Pyšek and Richardson 2007, Van Kleunen et al. 2010, Drenovsky et al. 2012, Gioria et al. 2023). Researchers commonly address this question by comparing the traits of invasive exotic species to those of both native and non-invasive exotic species within a taxon or an environment (Mathakutha et al. 2019). Examining invasive species alongside these two categories of non-invasive species helps distinguish traits associated with invasiveness from those that merely reflect introduction history (Van Kleunen et al. 2010, Moravcová et al. 2015). Throughout this study, we use the term *non-invasive species* to refer collectively to native species and exotic species that have not become invasive.

Plant traits can broadly be categorized into primary growth and secondary defense traits. Primary growth characteristics, such as specific leaf area, leaf nitrogen content, and leaf chlorophyll content, influence a plant’s ability to grow and reproduce. In contrast, secondary defense traits protect plants from environmental stressors, herbivores, and pathogens. These defenses can be further divided into physical defenses (e.g. leaf toughness and leaf thickness) and chemical defenses. Research on the contribution of primary growth traits to invasiveness is relatively well established: traits associated with rapid resource acquisition, growth, and reproduction consistently occur more strongly in invasive species than in non-invasive species (Van Kleunen et al. 2010, 2015). In contrast, studies of secondary plant defenses have generally found that defense investment differs less consistently between invasive and non-invasive species, suggesting that defense strategies may be more variable than growth-related traits (Lind and Parker 2010).

Although many studies have examined either primary growth characteristics or secondary plant defenses in invasive and non-invasive species, few have simultaneously evaluated both trait categories within the same comparative framework. Furthermore, studies that do investigate defenses tend to focus primarily on morphological traits, while secondary chemical defenses, especially widely distributed classes of specialized metabolites such as phenolics and terpenoids, have received comparatively little attention. These compounds are among the most widespread plant defense chemicals, reducing herbivory through toxic and deterrent effects while also protecting plants from environmental stress. Because the production of these metabolites requires substantial resources, variation in their investment may reflect trade-offs between defense and growth. As a result, it remains unclear whether coordinated differences in growth and defense strategies contribute to plant invasiveness. Consequently, a key gap remains in our ability to predict which introduced plant species are likely to become invasive.

This research aimed to address this gap by comparing primary growth and secondary defense traits of invasive and non-invasive *Lonicera* species. Non-invasive species included both native species and exotic species that have not become invasive, allowing us to distinguish traits associated with invasiveness from those associated simply with exotic origin. Specifically, we compared primary growth characteristics and secondary metabolite profiles across *Lonicera* species in the Arnold Arboretum of Harvard University to determine whether invasive species exhibit a distinct multivariate trait syndrome consistent with a growth-defense trade-off. Native *Lonicera* are widespread throughout New England, while several invasive exotic *Lonicera* species have become increasingly abundant across the region. In addition, the Arnold Arboretum of Harvard University houses an extensive collection of native, invasive exotic, and non-invasive exotic *Lonicera* species, providing an ideal system for comparing invasive species with a diverse set of non-invasive congeners.

Based on previous research on invasive plant traits, we predicted that invasive *Lonicera* species would exhibit greater investment in primary growth characteristics than non-invasive congeners (Van Kleunen et al. 2010, Schlaepfer et al. 2010). With respect to secondary defenses, the growth-differentiation balance hypothesis (Herms and Mattson 1992) predicts that plants under strong selection for rapid growth divert resources away from chemical defense, suggesting that invasive species may exhibit reduced investment in at least some chemical defense classes relative to non-invasive species. At the same time, invasive species may gain a competitive advantage by producing novel or particularly effective chemical defenses (Huberty et al. 2014). Therefore, we predicted that invasive *Lonicera* species would achieve higher growth potential through greater investment in primary growth traits while maintaining distinct, but not necessarily reduced, chemical defense profiles relative to non-invasive species.

## MATERIALS AND METHODS

This study focused on plant species in the genus *Lonicera* (Caprifoliaceae) and was conducted at the Arnold Arboretum of Harvard University, in Boston, Massachusetts, USA (42.298°N 71.123°W). The Arnold Arboretum maintains living collections of invasive and non-invasive *Lonicera* species, providing a shared study site for multivariate trait comparisons across plant guilds. We examined 14 *Lonicera* species in total: six invasive and eight non-invasive (Table S1). Of the non-invasive species, three were native and five were non-invasive exotic. Traits were assessed at two levels of organization (whole-plant and leaf-level) and spanned primary growth characteristics, physical defenses, and chemical defenses (Figure 1). Whole-plant traits were restricted to primary growth characteristics while leaf-level traits included all three trait categories.

**Figure 1.**
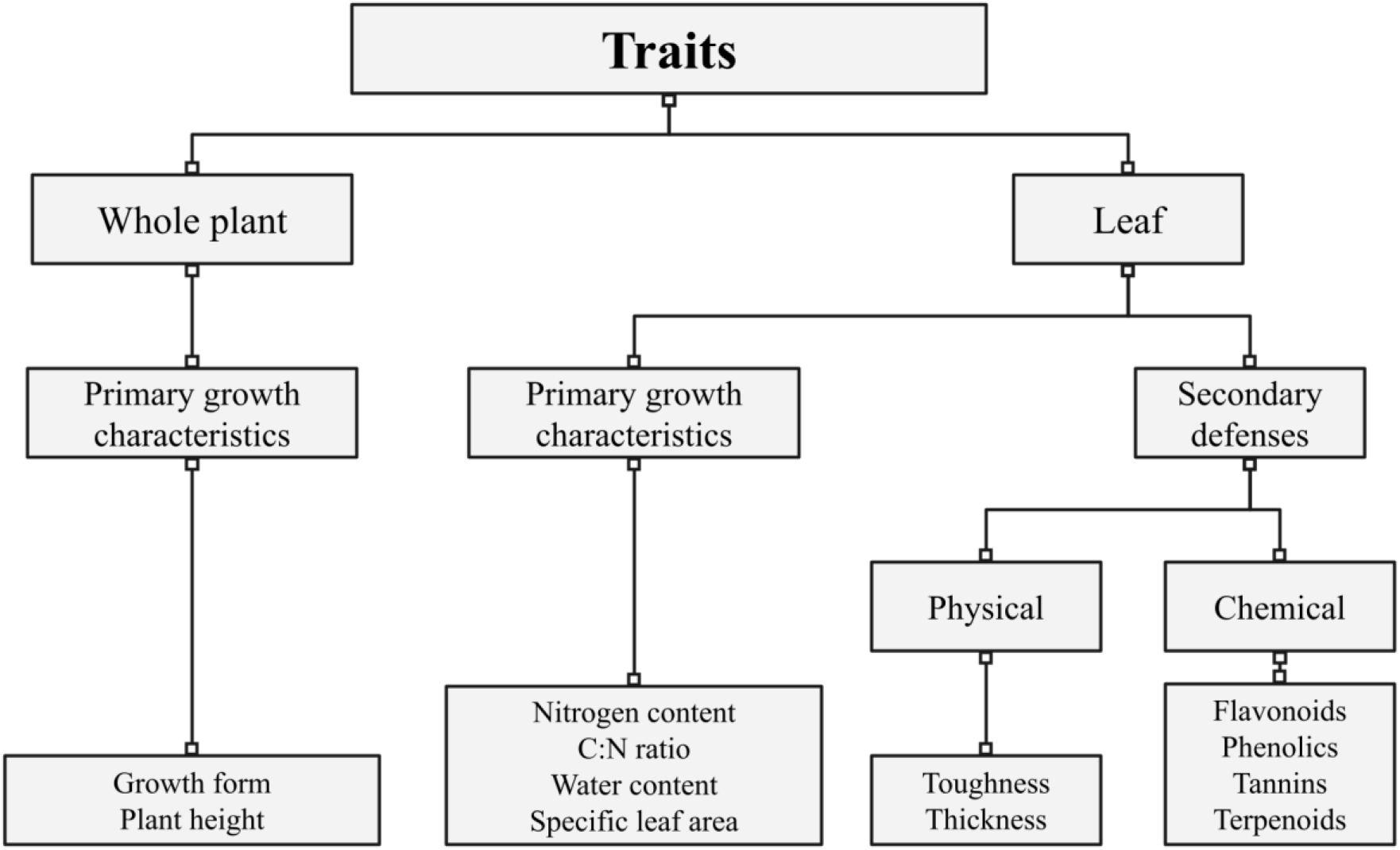
Flowchart of the traits examined in this study. The traits are first separated into whole-plant traits and leaf-level traits. The traits are then further divided into primary growth characteristics and (secondary) plant defenses. The plant defenses are divided into physical defenses and chemical defenses. We examined each of these traits on all 14 of the species included in the study.

### Whole-plant traits

We assessed two whole-plant primary growth characteristics: plant height and growth form. These traits were selected because they reflect environmental adaptation and reproductive ability, among other fitness-relevant attributes. Because the Arnold Arboretum only holds a select number of each species (occasionally only a single individual), we compiled species-level height and growth-form information from online botanical resources (Pérez-Harguindeguy et al. 2013, Native Plant Trust 2025). Sources consulted included *USDA Plant Database*, *Missouri Plant Finder*, and *Trees and Shrubs Online* to research our plants (Missouri Botanical Garden 2025, USDA Natural Resources Conservation Service 2025, Trees and Shrubs Online - Lonicera 2025). For primary growth form, each species was classified as a shrub or vine.

### Leaf-level traits

Leaf-level traits were chosen for their relevance to photosynthetic ability, herbivore resistance, and structural defenses (Funk and Throop 2010, Mathakutha et al. 2019). Field collections were conducted from July 7-11, 2025. For each species, five leaves were harvested to capture within-plant variation in light environment and position: one upper leaf (high up in the plant in the sun), a lower leaf (low to the ground), an inner leaf (closer to the trunk), an outer leaf (farther from the trunk in the sun), and a recently emerged (young) leaf. Harvested leaves were stored at 4°C until processing.

Each leaf was imaged using a flatbed scanner to obtain surface area and weighed to obtain wet mass. The leaf toughness was measured by using a digital penetrometer (FG3006, Shimpo Instruments) to puncture the lower half of each leaf three times halfway between the central vein and the margin and taking the median value recorded. We then measured the thickness of each leaf by using a micrometer in the left lower quadrant of the leaf, avoiding the central vein. The samples were then placed in a freeze dryer, where they were dried for 72 hours. Once the samples were dried, we calculated the combined dry mass of the five leaves, which was then averaged to get an average water content percent per leaf. The samples were then ground by hand using a mortar, pestle, and liquid nitrogen. Samples that needed further grinding were ground to a homogeneous fine powder using a Spex 8000M mill.

Leaf nitrogen and carbon percentages were measured using a FlashSmart 2000 CN analyzer. We also quantified total phenolics, condensed tannins, flavonoids, and terpenoids from freeze-dried ground leaf material using established colorimetric assays. All assays were performed on 25 mg subsamples. Total phenolics were estimated using the Folin-Ciocalteu assay (Ainsworth and Gillespie 2007), with extraction in 95% methanol for 48 hours in the dark and absorbance read at 765 nm. Condensed tannins and total flavonoids were quantified from a shared 100% methanol extract (five minute sonication, 30 minute dark incubation) using the vanillin–HCl and AlCl₃ assays, respectively, with absorbance read at 500 nm and 415 nm (Herald et al. 2014, Pękal and Pyrzyńska 2014). Terpenoids were estimated using the *p*-anisaldehyde assay with absorbance read at 600 nm (Le Bot et al. 2022). For all assays, extracts were clarified by centrifugation at 13,000 × *g* for five minutes, reactions were run against methanol blanks, and absorbance was measured by transferring 1500 µL of each reaction into a cuvette and analyzed with a single-slot visible light spectrometer (Vernier Software).

### Statistical analysis

All statistical analyses were performed using R version 4.4.1 “Race for Your Life” (R: The R Project for Statistical Computing 2024).

To compare overall traits among species, we ordinated the 14 *Lonicera* species in multivariate trait space using non-metric multidimensional scaling (NMDS) with the metaMDS function in the vegan package (Oksanen et al. 2025). The trait matrix comprised twelve traits per species: growth form, plant height, leaf nitrogen content, carbon-to-nitrogen ratio, flavonoids, phenolics, terpenoids, tannins, leaf toughness, leaf thickness, water content, and specific leaf area. Growth form, the only categorical trait, was coded numerically (shrub, vine) prior to ordination, and all traits were transformed onto a comparable uniform scale. The ordination was performed on a Bray-Curtis dissimilarity matrix in two dimensions (k = 2), using 20 random starts and up to 100 iterations to reduce the risk of convergence on a local minimum. We evaluated goodness of fit using the stress value, treating values below 0.2 as an adequate two-dimensional representation. To identify the traits most strongly associated with the ordination, we fit trait vectors to the NMDS using the envfit function with 999 permutations.

Our primary comparison was between invasive species (n = 6) and non-invasive species (n = 8), with native and non-invasive exotic species pooled in the latter group. This grouping addressed our central question of whether functional traits differed with invasion status. Group dispersion in the ordination was visualized using standard-deviation ellipses. We assessed differences in multivariate trait composition using permutational multivariate analysis of variance (PERMANOVA). We also explored three alternative grouping schemes: native versus invasive versus non-invasive exotic species; native vs “exotic” (comprising invasive and non-invasive exotic); and “competitive” vs “non-competitive” (in which “competitive” was comprised of native and invasive and “non-competitive” was comprised of non-invasive exotic). These additional comparisons were treated as exploratory rather than used to select the primary grouping.

We additionally tested each trait individually for differences among the two species guilds (invasive and non-invasive) using a simple linear model, with species guild as the sole predictor. Eleven traits were analyzed: plant height, leaf nitrogen content, carbon-to-nitrogen ratio, flavonoids, phenolics, terpenoids, tannins, water content, specific leaf area (SLA), leaf toughness, and leaf thickness. For leaf toughness and leaf thickness, the five leaves sampled per plant represented subsamples of a single individual rather than independent replicates; we therefore averaged these measurements to the species level prior to analysis, so that the species was the unit of replication for all traits (n = 14).

## RESULTS

This study examined 14 species of *Lonicera* plants, categorized into two functional groups: invasive and non-invasive species (which includes both native and non-invasive exotic species). The NMDS ordination (stress = 0.15) provided an adequate two-dimensional representation of differences in multivariate trait composition among species (Figure 2). A PERMANOVA indicated a significant difference in trait composition between invasive and non-invasive species (p = 0.048). Invasive species showed stronger correlations with plant height, water content, nitrogen content, flavonoids, and growth form, suggesting these species possess traits that favor rapid resource acquisition and competitive growth. These traits are characteristic of species that can quickly establish and spread in novel environments. In contrast, non-invasive species exhibited stronger associations with terpenoids, phenolics, tannins, leaf thickness, leaf toughness, and carbon-nitrogen ratio. This suite of traits points toward enhanced investment in structural and chemical defenses, suggesting that non-invasive species may prioritize defensive strategies over rapid resource acquisition.

**Figure 2.**
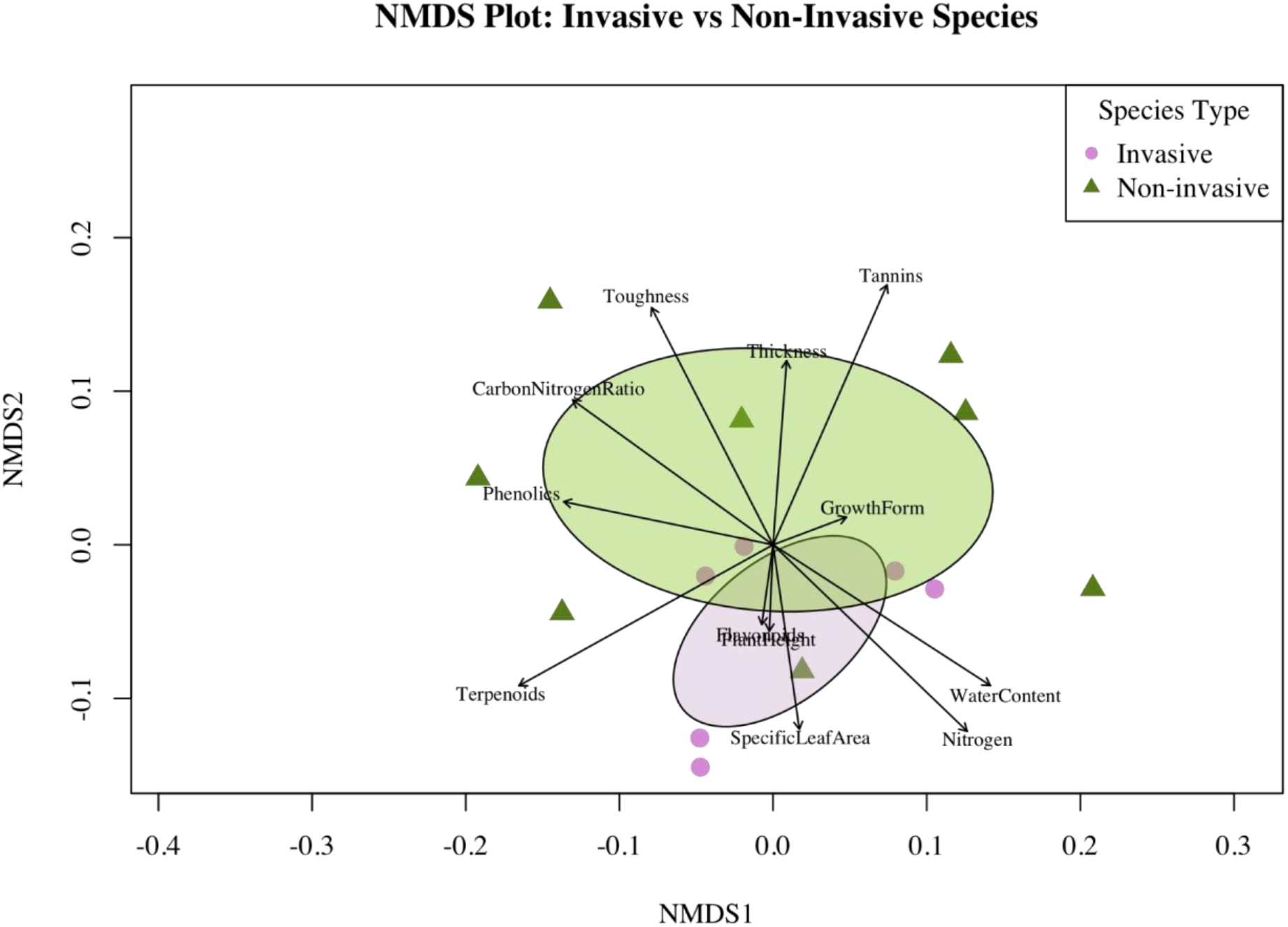
A non-metric multidimensional scaling (NMDS) is a two-dimensional representation of complex data, where points that are closer together are more similar to each other and are dissimilar to the other points. The NMDS plot demonstrates similarity across the two species guilds (invasive represented in purple circles and non-invasive represented in green triangles). Each point represents a single species. The ellipses highlight that there is clustering within each guild, indicating that the species within a guild are more similar to each other than to species in the other guild. The vectors displayed in the plot represent the direction and strength of each trait. A longer arrow indicates a stronger correlation between the ordination and the plant trait.

We investigated other species guilds by performing a PERMANOVA with four groups: native vs invasive vs non-invasive exotic, native vs “exotic”, “competitive” vs “non-competitive”, and invasive vs non-invasive. However, we recognized that the native vs “exotic” guild would not provide an accurate representation, as there were only three native species. Furthermore, the “competitive” vs “non-competitive” guild groups together native and invasive species, making it difficult to understand which functional traits relate specifically to invasiveness. Thus, we narrowed our comparison to the three-group species guild and the invasive vs non-invasive guild. PERMANOVA indicated a difference in multivariate trait composition between invasive and pooled non-invasive species (p = 0.048). Alternative groupings were examined as exploratory comparisons and were not used to establish the primary grouping.

We conducted a series of simple linear models comparing 11 leaf continuous traits between invasive and non-invasive plant species. Overall, we found two traits to differ significantly between the two groups: SLA and leaf thickness (Figure 3). For the whole plant traits, we detected no statistically significant differences between invasive and non-invasive plants. We observed no difference in plant height for the invasive and non-invasive species in this study (β_non-invasive_ = 7.75 ± 2.96, p = 0.075). Furthermore, of our primary growth characteristics, invasive species had significantly higher SLA than non-invasive species (β_non-invasive_ = 1.83 ± 0.29, p = 0.012), indicating that invasive plants tend to produce thinner, more efficient leaves per unit mass. However, for nitrogen content (β_non-invasive_ = 1.98 ± 0.27, p = 0.264), carbon-to-nitrogen ratio (β_non-invasive_ = 25.35 ± 2.93, p = 0.108), and average water content (β_non-invasive_ = 0.65 ± 0.03, p = 0.145), there was no difference between our species guilds.

**Figure 3.**
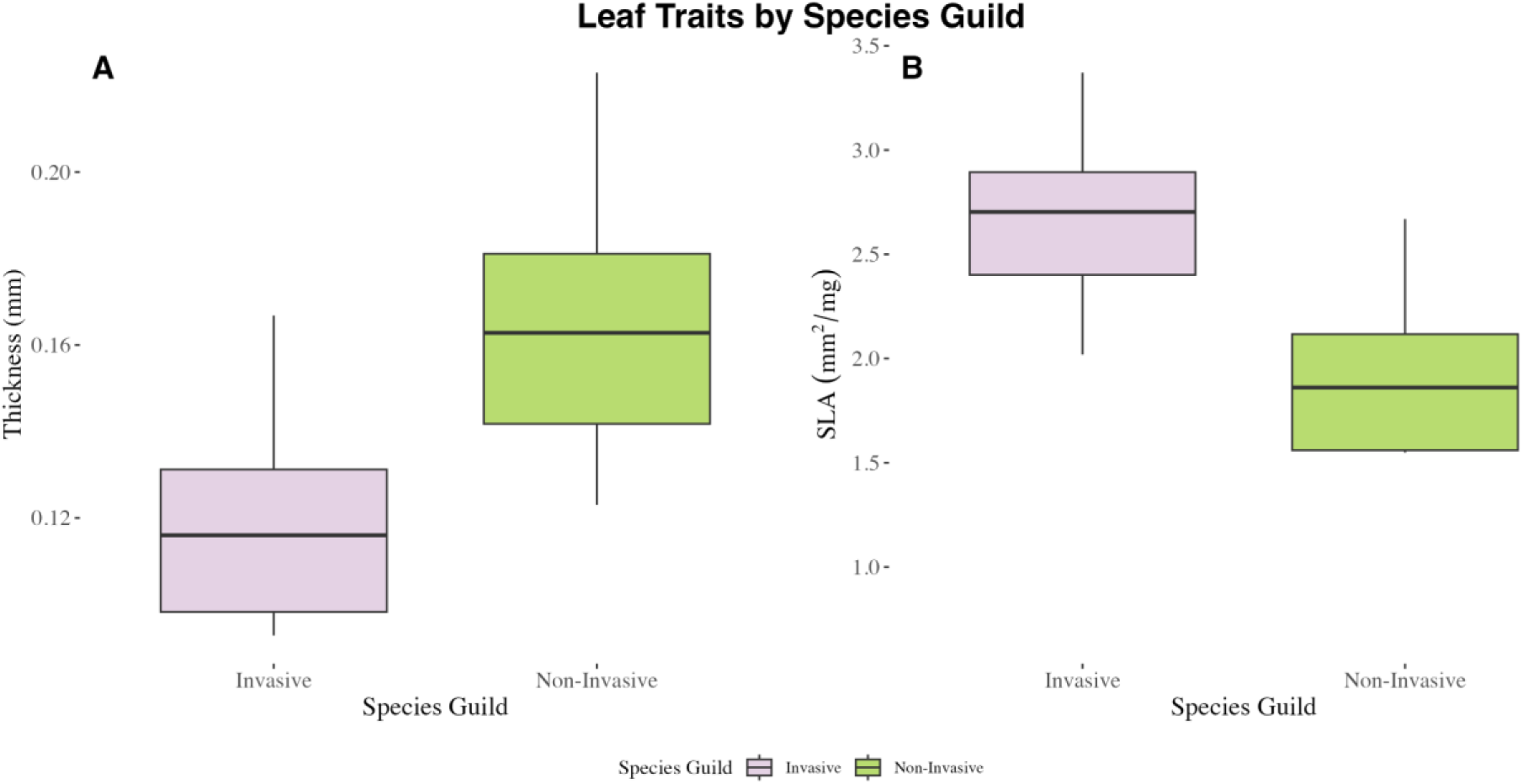
Boxplots for the significant results from the simple linear models for invasive (purple) and non-invasive (green) species. Panel A represents leaf thickness and panel B represents specific leaf area.

Looking at the secondary physical defenses, invasive species had significantly lower leaf thickness compared to non-invasive species (β_non-invasive_ = 0.17 ± 0.016, p = 0.017), but there was no difference in leaf toughness for invasive and non-invasive *Lonicera* species (β_non-invasive_ = 0.67 ± 0.08, p = 0.086). Finally, we found no difference in chemical defenses when comparing our two species guilds: flavonoids (β_non-invasive_ = 1.15 ± 0.20, p = 0.376), phenolics (β_non-invasive_= 0.51 ± 0.09, p = 0.375), terpenoids (β_non-invasive_ = 0.50 ± 0.17, p = 0.614), and tannins (β_non-invasive_ = 0.60 ± 0.15, p = 0.054).

## DISCUSSION

As invasive species become increasingly widespread, many studies have investigated whether invasive and non-invasive species differ in their functional traits. In this study, by grouping native and non-invasive exotic *Lonicera* species together as a single non-invasive category, we found that invasive *Lonicera* species differed from native and non-invasive exotic *Lonicera* in both primary growth and secondary defense traits.

Our results indicate that invasive species exhibited more traits associated with resource acquisition and rapid growth, while non-invasive species (comprising both native and non-invasive exotic species) exhibited more traits associated with plant defenses. The NMDS plot revealed separation between the two groups, with invasive species more strongly associated with plant height, growth form, nitrogen content, water content, SLA, and relative flavonoid concentration, all traits that largely reflect primary growth characteristics. In contrast, non-invasive species were more closely associated with leaf toughness, leaf thickness, carbon-to-nitrogen ratio, relative phenolics concentration, and relative tannin concentration, many of which are indicative of stronger structural and chemical defenses. These patterns were further supported by the simple linear model results; for instance, specific leaf area was significantly higher in invasive *Lonicera* species, a trait consistently linked to invasiveness in prior research (Mathakutha et al. 2019).

These findings are consistent with differences in growth and defense strategies between invasive and non-invasive plants. One possible explanation for the success of invasive species is the enemy release hypothesis (ERH), which proposes that invasive species experience reduced pressure from herbivores and pathogens in their introduced ranges (Keane and Crawley 2002). Reduced herbivore pressure may allow invasive species to maintain lower investment in certain defensive traits while exhibiting traits associated with rapid growth. Specifically for *Lonicera* species, Schweitzer and Larson (1999) found that an invasive *Lonicera* species exhibited greater morphological plasticity, potentially allowing it to adapt to varying environmental conditions. However, our results do not directly demonstrate resource allocation trade-offs between growth and defense. Although invasive *Lonicera* species exhibited higher specific leaf area and lower leaf thickness, we detected no significant differences in the chemical defense traits measured. These findings suggest that invasiveness in *Lonicera* may be associated with differences in growth-related and structural traits without necessarily involving reduced chemical defenses.

Furthermore, when we combined native and non-invasive exotic species into a single non-invasive group, we observed that these species were generally more defensive and less growth-oriented than invasive species. This demonstrates an observed trend in which non-invasive species are associated with a defense-oriented strategy. This pattern contrasts with some earlier studies that found trait divergence between natives and invasives specifically (Schweitzer and Larson 1999), but our study showed that the key functional difference is not between origin categories, but between invasive and non-invasive performance in the introduced environment.

In summary, our findings partially support our initial predictions: invasive *Lonicera* species exhibited higher specific leaf area and lower leaf thickness, traits consistent with differences in growth and structural defense strategies. However, we found no significant differences in the chemical defense traits measured. These findings suggest that invasiveness in *Lonicera* is associated with a combination of growth-related and structural traits, rather than a consistent reduction in chemical defenses. Our revised analytical framework also emphasizes the importance of grouping species by ecological impact rather than by nativity alone.

One limitation of this study is that trait comparisons were restricted to species within the genus Lonicera. Because trait patterns may vary across plant taxa, these findings may not generalize to other plant groups. Additionally, the number of individuals sampled was limited, as only one plant per species was available in the Arnold Arboretum. Increasing the number of individuals per species in future studies would allow for stronger comparisons while improving our confidence in the observed trait patterns. Finally, the set of primary growth and secondary defense traits examined in this study was limited by what could be measured within a single summer field season. Future research examining a broader range of growth and defense traits may provide a more comprehensive understanding of trait differences among invasive and non-invasive species.

Understanding which traits are associated with plant invasiveness is crucial for predicting future invasive events. Our results suggest that invasive Lonicera species exhibit distinct growth-related and structural traits compared with non-invasive species, particularly higher specific leaf area and lower leaf thickness. Identifying such trait patterns may improve our understanding of which introduced species are more likely to become invasive, potentially aiding the prevention of biological invasions.

## ACKNOWLEDGEMENTS

We thank the Arnold Arboretum of Harvard University for providing access to the plant specimens and resources necessary for this study. This research was supported by the DaRin Butz Foundation.

## APPENDIX

**Table S1.** Information on the 14 *Lonicera* species included in this study. We chose three native, six invasive, and five non-invasive exotic species based on what was available in the Arnold Arboretum (our field site). This table includes the species name, species group (native, invasive, or non-invasive exotic), the Arnold Arboretum accession number, and the native range of the species.

| Species name | Species group | Accession number | Native range or parentage |
|---|---|---|---|
| *Lonicera alpigena* (L.) | Non-invasive exotic | 1310-84*B | Central and southern Europe |
| *Lonicera* x *bella* (Zabel) | Invasive | 392-92*B | Cross between *L. morrowii* and *L. tatarica* |
| *Lonicera caerulea* (L.) | Native | 249-96*B | Holarctic |
| *Lonicera canadensis* (Bartram ex Marsh) | Native | 241-2017*D | Northeastern North America |
| *Lonicera floribunda* (Boiss. & Buhse) | Non-invasive exotic | 240-96*A | Central Asia |
| *Lonicera iberica* (M.Bieb.) | Non-invasive exotic | 486-95*A | Iran, Transcaucasus, Türkiye |
| *Lonicera japonica* (Thunb.) | Invasive | 2025-6*A | Japan, China, and Korea |
| *Lonicera maackii* ((Rupr.) Maxim.) | Invasive | 806-84*A | China |
| *Lonicera morrowii* (A.Gray) | Invasive | 368-95*C | Japan |
| *Lonicera nervosa* (Maxim.) | Non-invasive exotic | 95-81*D | China |
| *Lonicera sempervirens* (L.) | Native | 393-96*A | Eastern and southern United States |
| *Lonicera standishii* (Lindl. & Paxton) | Non-invasive exotic | 15118-3*A | China |
| *Lonicera tatarica* (L.) | Invasive | 299-78*B | Central Asia to south Russia |
| *Lonicera xylosteum* (L.) | Invasive | 856-84*A | Europe, Asia Minor, and W. Siberia |

## Notes

### Competing Interest Statement

The authors have declared no competing interest.

